# HortaCloud: An Open and Collaborative Platform for Whole Brain Neuronal Reconstructions

**DOI:** 10.1101/2025.03.13.642887

**Authors:** Konrad Rokicki, David Schauder, Donald J. Olbris, Cristian Goina, Jody Clements, Patrick Edson, Takashi Kawase, Robert Svirskas, Cameron Arshadi, David Feng, Jayaram Chandrashekar, Tiago A. Ferreira, MouseLight Project Team

## Abstract

HortaCloud is a cloud-based, open-source platform designed to facilitate the collaborative reconstruction of long-range projection neurons from whole-brain light microscopy data. By providing virtual environments directly within the cloud, it eliminates the need for costly and time-consuming data downloads, allowing researchers to work efficiently with terabyte-scale volumetric datasets. Standardization of computational resources in the cloud make deployment easier and more predictable. The pay-as-you-go cloud model reduces adoption barriers by eliminating upfront investments in expensive hardware. Finally, HortaCloud’s decentralized architecture enables global collaboration between researchers and between institutions.

## Introduction

State-of-the-art light microscopy techniques, combined with tissue clearing methods and computational pipelines, are successfully enabling mapping of complete neuronal projections at subcellular resolution in mammalian brains ^6,7,27^. However major challenges persist in big data handling, algorithmic accuracy, and scalable 3D visualizations ^20,21^. Despite recent advances in file formats ^19^, terabyte-scale light microscopy data sets are not easily accessible for interactive and collaborative neuron tracing.

As machine-learning based neuron tracing algorithms continue to improve ^8,16,28^, manual and semi-automated neuronal reconstruction paradigms (typically referred to as “tracing”) become even more pertinent for proofreading, verification, and ground truth generation. Tracing of sparse microscopy volumes is faster and more accurate when annotators take advantage of 3D Direct Volume Rendering (DVR) of image data ^27^. When image volume size exceeds GPU memory, DVR becomes challenging to implement. Out-of-core rendering strategies including octrees, multiscale representations (i.e. mipmaps), and file chunking can be implemented to allow visualization of multi-terabyte volumes. These strategies work well on a fast local area network (LAN) but when the data is moved to the cloud to improve collaboration, the latency for loading a chunk increases, degrading the user experience.

Overall, there are three possible deployment strategies for large-scale direct volume rendering software (Fig. 1): LL (local data, local rendering), CL (cloud data, local rendering), and CC (cloud data, cloud rendering). With *LL* systems (such as the original Horta ^14,27^ system and desktop tools like VVDViewer ^12^), the entire software stack is local and latencies are low. However, these types of systems do not support cross-institutional collaboration (or work-from-home, work-during-travel, etc.) because all data must be fully transferred to a local file store before it can be viewed. Although storage capacity has scaled to meet big data challenges, network transfer speed has remained largely fixed, especially for “last mile” delivery to end users. Transferring terabyte-scale data sets is slow and inconvenient, often requiring weeks of internet transfers or cumbersome physical transport of hard drives ^3^.

**Fig. 1.**
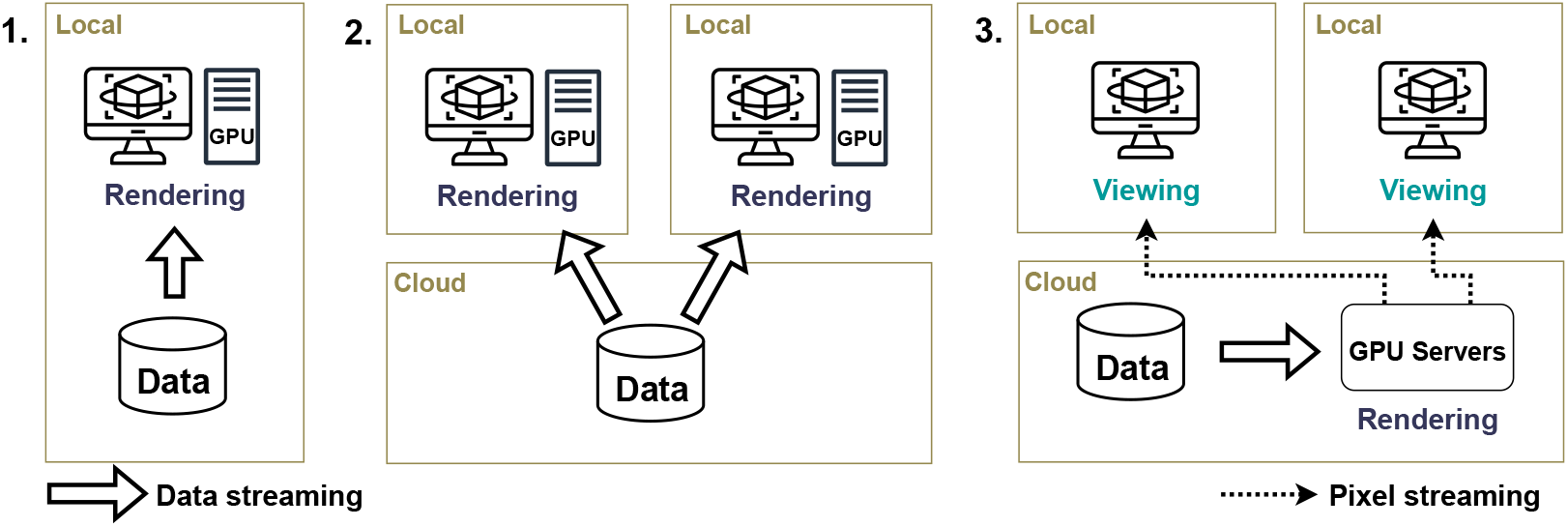
High-level architectural comparison. 1. Local data, local rendering (LL): Data and rendering share a local area network for fast connectivity; 2. Cloud data, local rendering (CL): Data is moved to the cloud, but must be streamed across the internet for local rendering; 3. Cloud data, cloud rendering (CC): Rendering is moved to the cloud, and only rendered pixels are streamed to the clients.

In *CL* systems, data is moved to the cloud to improve accessibility and support cross-institute collaboration. However, because the rendering is performed locally, the data need to be downloaded across the internet before they can be rendered. This process can introduce significant delays, especially for clients that are geographically distant and those with slower internet connections. CL systems often run within web browsers to take advantage of the web’s advantages in application distribution and accessibility. However, web browsers have memory limits at multiple levels—such as browser limit, single tab restrictions, and limitations within WebGL and WASM—and cannot easily make use of large amounts of system memory. In addition, historically web browsers have not had access to robust graphics primitives or libraries. This is starting to change with recent advances in WebGPU and WebAssembly but web-based CL systems remain an active area of research ^25^. One example of a popular CL system is Neuroglancer, a web-based visualization tool capable of handling large images. Although it includes an experimental DVR feature, it is primarily optimized for 2D cross-sectional views. Other web-based platforms ^1,22^ for large microscopy image visualization and neuron annotation are optimized for dense reconstructions in electron microscopy (EM) images and face limitations when applied to large-scale volume rendering of sparse light microscopy data. Although they allow users to collaboratively annotate neuron skeletons in large 3D image stacks, the images stacks are only rendered in cross-sectional views.

These challenges with LL and CL systems exist even outside of the scientific visualization domain and have prompted many big data practitioners to adopt the fully cloud-based paradigm (*CC*) where computation is brought to data, instead of moving data to compute facilities ^17^. To support interactive 3D graphics, cloud-based Virtual Desktop Infrastructure (VDI) solutions render users’ desktops in the cloud, and stream the rendered pixels back to a lightweight browser-based client. This approach is gaining tremendous popularity for gaming ^2^, and has many benefits besides faster data loading latency. Processing and visualizing large 3D datasets often demand specialized hardware, which can be challenging for many labs to acquire and manage. ^9,15,24^. By contrast, cloud systems can be rented on an hourly basis, eliminating the need for upfront capital investments. Second, deploying complex 3D rendering applications on local systems is complicated by the fact that hardware and OS configurations can vary widely across institutions. Cloud systems provide a way to standardize the environment across deployments and greatly reduce code maintenance overhead. Finally, deployment to cloud systems can be effectively standardized and automated through the use of Infrastructure as Code (IaC) provisioning.

We encountered such challenges with the Horta ^14^ neuron tracing software, an LL system which was used by the MouseLight Project Team to reconstruct over 1000 neurons in the mouse brain ^27^. Horta is a client-server system that was deployed on our internal servers and used by a team of annotators for many years. However, deploying Horta at other institutions revealed several limitations. Although deployment of the Horta system was streamlined using Docker Swarm, managing the deployment still required a high level of DevOps expertise. Collaborator institutions ran into many issues attempting to deploy Horta on their own infrastructure, where differences in operating systems, path mounting, hard disk sizes, and other institution-specific computing practices created subtle problems which took long to debug. In addition, some institutions were put off by the expenses of acquiring servers and high-end desktop workstations for their annotators.

## Implementation

To overcome these challenges, we created HortaCloud by transitioning the entire Horta infrastructure to the cloud (Fig. 2) using Amazon Web Services (AWS). The data was uploaded to Amazon S3, a distributed key value store. The backend services were deployed on an Amazon Elastic Compute Cloud (EC2) instance. The client was deployed via Amazon AppStream 2.0, a VDI which automatically provisions GPU-equipped virtual machines as users log in. We also created a portal website which lets administrators manage user accounts, and allows users to launch new application sessions.

**Fig. 2.**
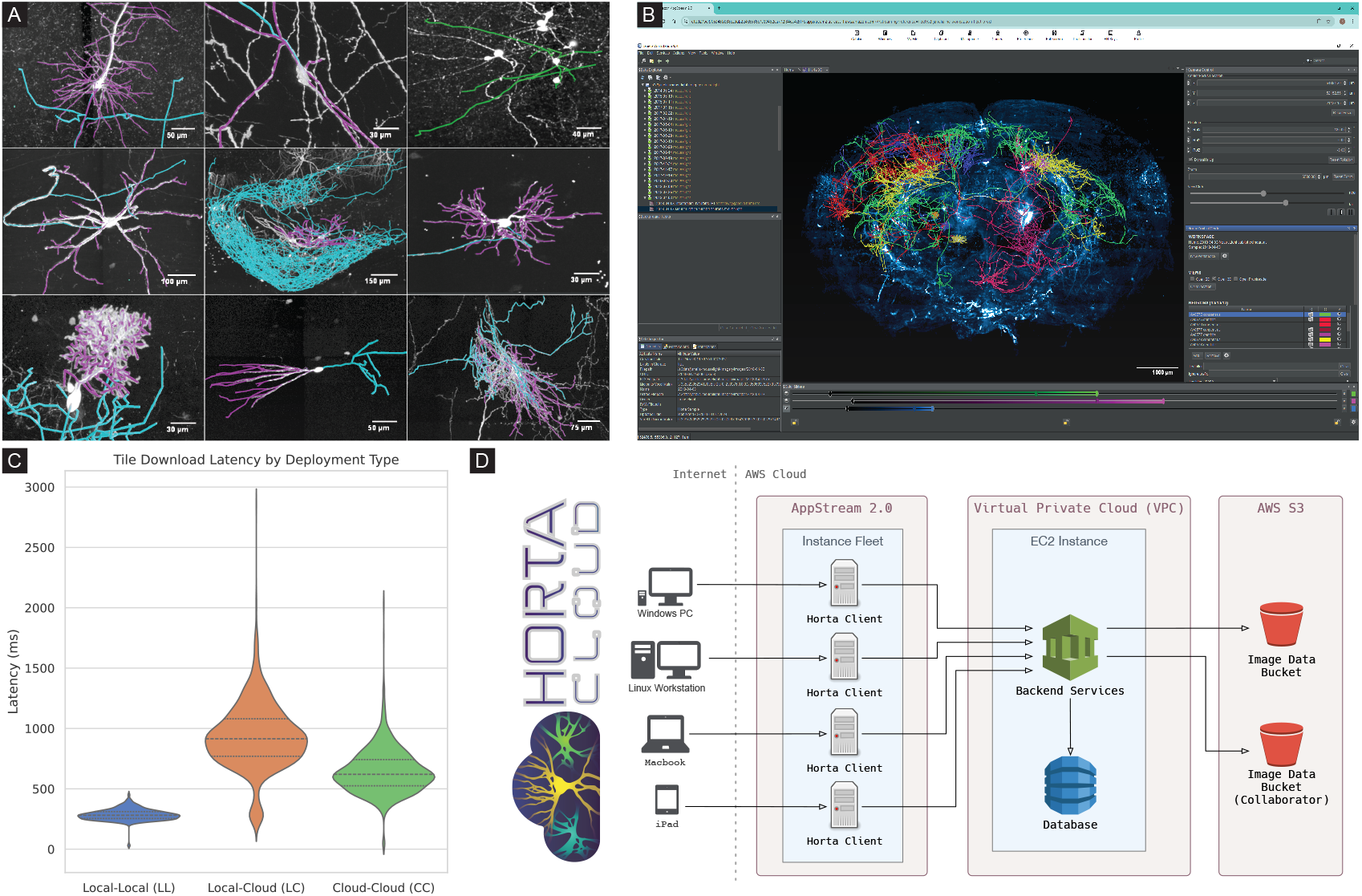
Overview of HortaCloud. (A) Diversity of neuronal morphologies found in the adult mouse brain (detailed in Supplemental Section II). Axonal arbors feature complex topologies that can only be reconstructed with fidelity by means of responsive and interactive rendering of high resolution 3D imagery. Note that in most cases axonal arbors span dozens of centimeters beyond the displayed fields of view. (B) A screenshot of HortaCloud running in a web browser. A MouseLight sample is loaded, with published neurons displayed. The left-hand panel shows other datasets which can be loaded. The right-hand panel shows details about the current sample and workspace, including individual neurons. Color sliders at the bottom allow fine adjustment of the image rendering, including spectral unmixing operations. (C) Comparison of KTX tile loading latencies across the tested deployment types. The 2023-09-01 MouseLight sample was loaded into Horta and the view was automatically moved across the sample using the Neuro Cam feature, forcing many tiles to be download. We collected the latency of 1500 tile downloads for each deployment. (D) A high-level architectural overview of the HortaCloud system. Clients connect to AppStream and use shared backend services and database running on AWS EC2. Data is served from AWS S3 buckets.

HortaCloud is deployed at the institutional level, with each deployment instance supporting a team of annotators working collaboratively. Each instance is backed by a shared database and suite of RESTful services deployed on EC2, which support collaboration features such as data sharing. The system automatically scales with usage by creating a new virtual desktop for each user. The Allen Institute for Neural Dynamics (AIND) hosts an average of 40 concurrent users on their HortaCloud instance.

Importantly, Horta was optimized for both performance and reconstruction accuracy ^5^. Key functionality includes:

- Extensive proofreading controls
- Support for up to a million skeletonized annotations per workspace
- Concurrent, multi-user tracing, in which access to shared data is managed by a permissions framework
- Dynamic controls for spectral unmixing, in which background autofluorescence acquired as a separate channel on the microscope can be subtracted on the fly from the channel being traced, thus enhancing the signal to noise ratio along neurites
- Manual and semi-automated tracing workflows

We modified Horta in several ways to better integrate it with the cloud environment. The existing Horta data service was only able to read from local storage and Network File System (NFS) mounts. We added the capability to read from any S3 bucket, to give users the ability to load any data residing in the AWS cloud. We also implemented support for the OME-Zarr next generation file format ^19^, the emerging standard for storing biomedical imaging data in the cloud. Horta also supports its own octree file format based on KTX (Khronos Texture) that can be streamed directly into GPU memory. Next, we created a Data Importer tool which can be used to convert various image formats to KTX octrees. Finally, we integrated the Horta desktop client with the AppStream service, so that it is launched when the AppStream session starts, and the user is automatically logged in.

More details about the implementation can be found in the Supplementary Information.

## Results and Discussion

HortaCloud is a powerful computational environment that can accelerate scientific efforts such as whole brain neuron tracing and other collaborative large volume annotation projects. It has been used for whole brain neuron tracing in mammals ^4,5,27^ and visualization of their microvasculature ^11^. The HortaCloud application is accessed through any modern web browser and provides instant access to hundreds of image datasets and thousands of neuron tracings. After logging into HortaCloud, users are presented with a remote desktop interface running Horta. Users can add any S3 bucket, public or private, and load compatible images into the viewer for viewing or tracing. We used the Data Importer to convert an fMOST dataset ^7^ from the Brain Image Library ^13^ into KTX format and load it in HortaCloud. AIND loaded their public ExA-SPIM data ^6^ (OME-Zarr formatted) into HortaCloud, as well as other fMOST imagery.

The tracing environment is pre-populated with most of the imagery used to create the collection of single neuron reconstructions available in the MouseLight Neuron Browser, along with their traced neuron skeletons. These datasets feature high-fidelity reconstructions of axonal projections, each traced twice by different annotators ^5,23^. By making the imagery and the reconstructions available, we hope to facilitate analysis and validation of reconstruction quality at the whole-brain scale. Images can be opened for viewing in Horta 2D or 3D views for verification and further tracing annotation can be performed. Horta allows users to save and share “deep-links” which include the 3D location in the sample, camera orientation, zoom level, and other contextual information. These deep-links enable seamless navigation to specific points in the sample, making it easy to revisit annotations, and share views of interesting brain locations.

An important factor for tracing efficiency is the latency of tile loading when following neurons through the volume. In mammals, neuronal axons typically span a substantial portion of the brain volume, requiring constant tile loading as the user moves along the neuron. We compared the latency of tile loading in each of the three deployment strategies (Fig. 2C). We found that the LL instance offered the fastest tile loading with the lowest amount of variability. This was expected as our local network and storage systems are highly optimized for both latency and throughput. With data in the cloud, the LC deployment had nearly twice as much latency for KTX tile retrieval as the CC deployment. Our CC deployment is slower than LL, because we use standard S3 storage which is not optimized for access speed or throughput. However, in our experience it is still fast enough for effective tracing. It is possible to use high-performance storage in the cloud (e.g. S3 Express One Zone) to make the performance comparable to optimized local systems or better, but the additional cost does not seem justified for our use cases.

Ongoing operational costs are a common concern with cloud deployments. Over the past several years, we have found the cost for our HortaCloud instance to be comparable to our bare-metal deployments, even without accounting for the total cost of ownership (TCO) including hardware refresh, system administration, and desktop support. In addition, cloud deployments do not require upfront investment and provide more flexibility when usage is less than full-time. We make updated cost estimates available on the HortaCloud website via the AWS Cost Calculator. As of 2024, the TCO of a HortaCloud deployment is approximately $748 per month for a single full-time user, and $178 per month for each additional user (the complete estimates are included as Supplemental Files). These costs are estimated using on-demand EC2 instances and can be reduced by using a pre-paid plan. We were able to take advantage of the AWS Open Data Sponsorship program to publish data in public S3 buckets free of charge, which greatly reduces our overall cloud costs.

Finally, we are launching HortaCloud Community as an open source project that accepts contributions via GitHub. In this way, the software can continue to evolve to meet new needs, and the scientific community will benefit from institutional engagement in the software’s upkeep. The HortaCloud Community can be used as a resource to discuss issues, propose solutions, and route information to developers when fixes are necessary. In addition to detailed documentation, we have also released step-by-step protocols that guide new users into technical workflows ^23^.

## Supporting information

AWS Cost Estimate (1 user)

AWS Cost Estimate (2 users)

Tile Loading Latency Data

## Data Availability

MouseLight imagery ^27^ and associated un-transformed reconstructions are accessible through all HortaCloud instances. The data is hosted on AWS Open Data in the s3://janelia-mouselight-imagery S3 bucket, under a CC BY 4.0 license. AIND makes its ExA-SPIM data available in OME-Zarr format in the s3://aind-open-data ^6^ S3 bucket, also under a CC BY 4.0 license. Other data accessible through HortaCloud instances may be available under different licenses and terms-of-use.

The latency data underlying Fig 2C is included in the Supplemental Files (`latencies.csv`).

## Code Availability

The complete source code for HortaCloud is available in the repositories listed below. Documentation and other information is hosted at https://hortacloud.org.

- Automated Deployment
  - Source code repository: https://github.com/JaneliaSciComp/hortacloud
  - Archived code at time of publication: https://doi.org/10.5281/zenodo.15002731
- Janelia Workstation (including Horta client)
  - Source code repository: https://github.com/JaneliaSciComp/workstation
  - Archived code at time of publication: https://doi.org/10.5281/zenodo.15002705
- JACS Configuration Management
  - Source code repository: https://github.com/JaneliaSciComp/jacs-cm
  - Archived code at time of publication: https://doi.org/10.5281/zenodo.15002726
- JADE Storage Engine
  - Source code repository: https://github.com/JaneliaSciComp/jacs-storage
  - Archived code at time of publication: https://doi.org/10.5281/zenodo.15014843
- JACS Compute Services
  - Source code repository: https://github.com/JaneliaSciComp/jacs-compute
  - Archived code at time of publication: https://doi.org/10.5281/zenodo.14968242
- JACS Domain Model
  - Source code repository: https://github.com/JaneliaSciComp/jacs-model
  - Archived code at time of publication: https://doi.org/10.5281/zenodo.14932938
- JACS Message Broker
  - Source code repository: https://github.com/JaneliaSciComp/jacs-messaging
  - Archived code at time of publication: https://doi.org/10.5281/zenodo.15014855
- Website and documentation
  - Source code repository: https://github.com/JaneliaSciComp/hortacloud-website
  - Archived code at time of publication: https://doi.org/10.5281/zenodo.15014859
- Data importer
  - Source code repository: https://github.com/JaneliaSciComp/hortacloud-importer
  - Archived code at time of publication: https://doi.org/10.5281/zenodo.7586244

## Author Contributions

D. Schauder, D.J. Olbris, C. Goina, K. Rokicki, J. Clements, P. Edson, T. Kawase, and C. Arshadi implemented the software. R. Svirskas uploaded and managed the data on AWS. T.A. Ferreira, J. Chandrashekar, K. Rokicki, and the MouseLight steering committee conceptualized the project. D. Feng contributed technical advice. The MouseLight Project Team and Allen Institute for Neural Dynamics were the primary users of the software and contributed bug reports and ideas for features. K. Rokicki, T.A. Ferreira, and D. Schauder wrote the manuscript. All authors reviewed and approved the final version of the manuscript.

The MouseLight Project Team during this time consisted of: Andrew Recknagel, Cameron Arshadi, Emily Tenshaw, Jayaram Chandrashekar, Mary Lay, Monet Weldon, Patrick Edson (Leap Scientific), and Tiago A. Ferreira. Joshua T. Dudman, Adam Hantman, Scott Sternson, Karel Svoboda, Nelson Spruston, and Wyatt Korff were part of the MouseLight steering committee.

## Acknowledgments

We thank Stephan Preibisch and John Bogovic for thoughtful feedback about the draft manuscript; Alyssa Stark, Emily Tenshaw, and Mary Lay for collecting usage data; Janelia Scientific Computing, especially Adam Taylor and Ben Arthur for help with data processing; Ken Carlile and Scientific Computing Systems for help with data handling; Christopher Bruns for foundational software development on the Horta tool; Sean Murphy, Todd Safford, Eric Trautman, and Les Foster for early work on the Janelia Workstation framework and Horta tool; Brie Yarbrough for creating the HortaCloud logo. We are grateful to all those who provided bug reports and suggestions on the software.

This work was supported by the Howard Hughes Medical Institute, Janelia’s Open Science Software Initiative (OSSI), the MouseLight Project Team, and the AWS Open Data Sponsorship Program. We used diagram images from draw.io and flaticon.com. This article is subject to HHMI’s Open Access to Publications policy. HHMI lab heads and Project Team Leaders have previously granted a nonexclusive CC BY 4.0 license to the public and a sublicensable license to HHMI in their research articles. Pursuant to those licenses, the author-accepted manuscript of this article can be made freely available under a CC BY 4.0 license immediately upon publication.

## Supplementary Information

### I. Software Architecture

HortaCloud is a cloud deployment of Horta, a neuron tracing application built on top of the Janelia Workstation platform. This manuscript is focused on the HortaCloud deployment and its cloud-based architecture. However, summaries of the Workstation platform and Horta application are provided here for context.

#### I.1 Janelia Workstation

Janelia Workstation ^14^ is a platform for building integrated data management environments for neuroscience research. Specifically, it was built to support large scale confocal image processing for the FlyLight project at Janelia Research Campus ^10^. Later, the same platform was used as a basis for Horta, supporting the MouseLight Project Team. The Janelia Workstation platform code is publicly available and licensed for reuse under the BSD 3-Clause open source license.

The Janelia Workstation platform includes a NetBeans-based Java thick client which supports cross-platform installation, remote auto-updates, dockable UI components, a menu-based action system, key bindings, user authentication and permissions, remote domain objects with faceted full text search, object hierarchies and annotations, and other foundational features for building data management applications.

From an architectural perspective, the Janelia Workstation is a client-server application with a microservice back-end (Fig. 3) supporting state management and remote data access. The micro services are fronted with an API gateway, and include: JACS sync web services for authentication, user management, and metadata access, JACS async web services which process data in the background either locally or on a compute cluster, JACS messaging service which relies on RabbitMQ to route asynchronous messages between clients and services, JADE web services for remote access to data storage which scale horizontally with multiple worker nodes, a full text indexing service implemented with SOLR, and a MongoDB database cluster for scalable data management.

**Fig. 3.**
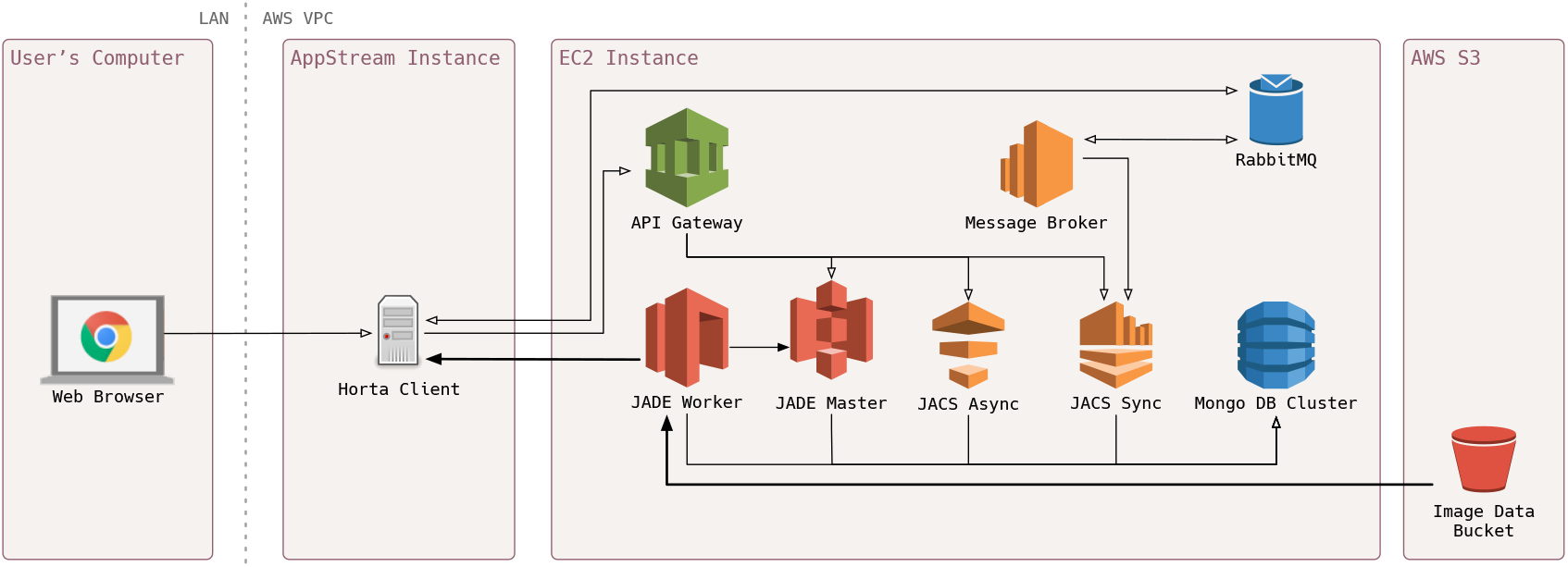
Microservice Architecture. Deployment diagram showing the microservices comprising the JACS backend to the Janelia Workstation. This includes the API Gateway which provides routing and SSL termination, the JADE system for storage access, the JACS system for metadata access and asynchronous compute, the MongoDB cluster, RabbitMQ instance for messaging, and the message broker. Microservices are deployed using Docker Swarm and can be scaled across multiple EC2 instances. The AppStream instance runs the Horta client and serves it to the user’s browser.

The microservice back-end is containerized using Docker and deployed across multiple servers using Docker Swarm. An associated ELK stack (Elasticsearch, Kibana, and Logstash) provides server log indexing and searching across all microservices.

#### I.2 Horta

The Horta application consists of interconnected modules built on the Workstation platform. These modules include a multi-scale, out-of-core 3D renderer/viewer, a 2D tile-based viewer, and ergonomic neuron-tracing tools designed for efficient annotation. Multi-level resolution tiles are dynamically queued and loaded into the viewer based on the user’s zoom-level and camera focus, allowing mesoscale visualization at various levels of magnification. Horta also optimizes data management for images through parallel loading and multi-resolution pyramids, enabling fast visualization of multi-terabyte datasets. The 3D viewer supports free rotation around the sample as well as panning and zooming the view using mouse controls or precise location coordinates to quickly navigate to any location in the sample.

Core features for rapidly annotating neurons include operations such as appending to a neuron, splitting, merging, re-rooting, and moving neuron vertices. Annotators can add and review notes on each vertex or branch and group neurons to toggle visibility or assign ownership. Neuron information such as number of vertices and total length can be generated, along with a neuron review feature that creates a dendrogram view and allows users to review a neuron by automatically panning the camera along its branches.

Horta enables real-time collaborative tracing for multiple annotators working on the same sample. Its permissions model prevents users from accidentally modifying each other’s annotations, while asynchronous messaging using RabbitMQ publish/subscribe framework ensures users can view and stay updated on one another’s changes in real time.

For machine-learning assisted workflows, Horta supports the import of hundreds of thousands of ML-generated neuron fragments into a single sample. It uses MongoDB GridFS and a spatial index to dynamically load only those fragments in a radius around the user viewport. These fragments can then be merged by annotators at key crossover points or reviewed to greatly accelerate neuron reconstruction.

Finally, it supports data import/export with interoperable file formats such as SWC (validated ^18^ against the latest specification) and facilitates movie generation by means of a built-in movie recorder. It also supports overlaying meshes (Wavefront OBJ) of neuropil labels (such as compartments of the Allen Mouse Brain Common Coordinate Framework ^26^) into the sample for visualizing neuron relationships to different brain regions.

#### I.3 HortaCloud

The architectural keystone of HortaCloud is the ability to deploy both the clients and servers on a single Virtual Private Cloud (VPC). This allows the large image data to stay in the cloud where it is streamed and rendered, while users can view a high-quality video stream of the interactive desktop in any modern web browser.

HortaCloud leverages the AppStream 2.0 for rendering and streaming to the client. AppStream provides a VDI solution where the Horta client can run on a secure private network with GPU rendering capability. In particular, AppStream is based on the NICE DCV protocol which uses library interposing to intercept OpenGL calls and redirect them to a virtual machine for rendering. Any 3D windows are efficiently rendered on the GPU, and the resulting pixels are compressed using the H.264 video codec, and streamed directly back to the client to be transparently recomposed with the rest of the desktop. When streaming, the parts of the desktop which are changing are sent over the network, reducing bandwidth requirements. We use stream.graphics.g4dn.xlarge instances.

Back-end microservices are deployed to one or more EC2 instances and orchestrated using Docker Swarm. The EC2 instances are located in the same VPC as the AppStream client, ensuring that all data is transferred securely, and service endpoints are not open to Internet traffic. We use a r5n.2xlarge instance to run all of the microservices, including the MongoDB cluster. Persistent data from the MongoDB cluster is backed up on a nightly basis to S3.

Image data is hosted on S3, a scalable file service with optional cross-region replication. S3 has high durability and is configurable for different usage patterns (e.g. intermittent versus constant), and is cost effective for large image storage.

To simplify accessibility, we integrated Horta’s user authentication system with AWS Cognito. User accounts are created with a web portal which synchronizes them across Cognito and the JACS back-end with an AWS Lambda function. The web portal allows administrative users to delete users or temporarily block their access. When a user logs into the web portal and opens AppStream, they are automatically logged into the Horta application.

The entire HortaCloud infrastructure is generated using the AWS CDK framework. CDK allows us to programmatically define the virtualized cloud components and dependencies between them, and greatly simplifies the deployment process while reducing DevOps toil and application administration overhead.

### II. List of Morphological Exemplars (Relates to Figure 2)

Neurons depicted in Figure 2A were collected from MouseLight datasets. Reconstructed dendrites are highlighted in magenta, axons in cyan, and unclassified neurites in green. Screenshots depict a small field of view and do not include the complete axonal arbor, which typically extends several centimeters away from the neuron’s cell body. Exemplars are located on the following brain regions (left to right, top to bottom): Primary motor cortex [MouseLight sample (MLS) 2023-02-02]; Intermediate reticular nucleus [MouseLight neuron (MLN) AA0434]; Hypothalamus [MLS 2017-02-22]; Superior central nucleus raphe [MLN AA1589]; Dentate gyrus (polymorph layer) [MLN AA1393]; Thalamus [MLS 2017-06-10]; Flocculus [MLN AA0963]; Dentate gyrus (molecular layer) [MLN AA0166]; Primary visual area [MLN AA0028].

