## Supplementary material for "HortaCloud: An Open and Collaborative Platform for Whole Brain Neuronal Reconstructions": AWS Cost Estimate (1 user)

Contact your AWS representative: Contact Sales

**Export Date:** 12/18/2024 **Language:** English

Estimate url

### Estimate summary

| Upfront cost | Monthly cost | Total 12 months cost |
|---|---|---|
| 0.00 USD | 748.43 USD | 8,981.16 USD<br>Includes upfront cost |

### Detailed Estimate

| Name | Group | Region | Upfront cost | Monthly cost |
|---|---|---|---|---|
| Amazon Simple Storage Service (S3) | - | US East (N. Virginia) | 0.00 USD | 76.06 USD |

**Status** -

**Description:** Private sample storage

**Config summary** S3 Standard storage (3 TB per month), PUT, COPY, POST, LIST requests to S3 Standard (1000000), GET, SELECT, and all other requests from S3 Standard (1000000) DT Inbound: Internet (12 TB per month), DT Outbound: Not selected (0 TB per month)

| Name | Group | Region | Upfront cost | Monthly cost |
|---|---|---|---|---|
| Amazon AppStream 2.0 | - | US East (N. Virginia) | 0.00 USD | 182.54 USD |

**Status** -

**Description:** -

**Config summary** License model (Commercial license included), Instance type (stream.graphics.g4dn.xlarge), Operating system (Windows), Days in week (5), Peak duration (hours) per day (8), Average off-peak concurrent users per hour (0), Average peak concurrent users per hour (1), Days in weekend (0), Peak duration (hours) per day (0), Average off-peak concurrent users per hour (0), Average peak concurrent users per hour (0)

| Name | Group | Region | Upfront cost | Monthly cost |
|---|---|---|---|---|
| Amazon EC2 | - | US East (N. Virginia) | 0.00 USD | 456.08 USD |

**Status** -

**Description:** Backend services

**Config summary** Tenancy (Shared Instances), Operating system (Linux), Workload (Consistent, Number of instances: 1), Advance EC2 instance (r5n.2xlarge), Pricing strategy (On-Demand Utilization: 100 %Utilized/Month), Enable monitoring (disabled), EBS Storage amount (210 GB), DT Inbound: Not selected (0 TB per month), DT Outbound: Not selected (0 TB per month), DT Intra-Region: (0 TB per month)

| Name | Group | Region | Upfront cost | Monthly cost |
|---|---|---|---|---|
| Amazon Virtual Private Cloud (VPC) | - | US East (N. Virginia) | 0.00 USD | 33.75 USD |

**Status** -

**Description:** VPC

**Config summary** Number of NAT Gateways (1) DT Inbound: Not selected (0 TB per month), DT Outbound: Internet (10 GB per month), DT Intra-Region: (0 TB per month)

**Acknowledgement**
AWS Pricing Calculator provides only an estimate of your AWS fees and doesn't include any taxes that might apply. Your actual fees depend on a variety of factors, including your actual usage of AWS services. Learn more
